# Low levels of transposable element activity in *Drosophila mauritiana*: causes and consequences

**DOI:** 10.1101/018218

**Authors:** Robert Kofler, Christian Schlötterer

## Abstract

Transposable elements (TEs) are major drivers of genomic and phenotypic evolution, yet many questions about their biology remain poorly understood. Here, we compare TE abundance between populations of the two sister species *D. mauritiana* und *D. simulans* and relate it to the more distantly related *D. melanogaster*. The low population frequency of most TE insertions in *D. melanogaster* and *D. simulans* has been a key feature of several models of TE evolution. In *D. mauritiana*, however, the majority of TE insertions are fixed (66%). We attribute this to a lower transposition activity of up to 47 TE families in *D. mauritiana*, rather than stronger purifying selection. Only three families, including the extensively studied *Mariner,* may have a higher activity in *D. mauritiana*. This remarkable difference in TE activity between two recently diverged *Drosophila* species (≈ 250,000 years), also supports the hypothesis that TE copy numbers in *Drosophila* may not reflect a stable equilibrium where the rate of TE gains equals the rate of TE losses by negative selection. We propose that the transposition rate heterogeneity results from the contrasting ecology of the two species: the extent of vertical extinction of TE families and horizontal acquisition of active TE copies may be very different between the colonizing *D. simulans* and the island endemic *D. mauritiana*. Our findings provide novel insights in the evolution of TEs in *Drosophila* and suggest that the ecology of the host species could be a major, yet underappreciated, factor governing the evolutionary dynamics of TEs.

## Significance statement

Transposable elements (TE) are stretches of DNA that selfishly propagate within genomes. By comparing TE abundance between closely related *Drosophila* species we found that TE activity is significantly lower in an island endemic species than in worldwide distributed species. We propose that geographic isolation on islands could lead to fewer opportunities for acquiring active TEs by horizontal transfer as well as an accelerated loss of functional TEs. Our results suggest that ecology of host species may be a powerful factor which modulates the activity of multiple TE families even over very short evolutionary time scales.

## Introduction

Transposable elements (TEs) are stretches of DNA that selfishly spread within genomes, even to the detriment of the host (Hickey, 1982). Insertions of TEs in host genomes may have a significant impact on phenotypes, including diverse phenomena such as variation of quantitative traits (Mackay et al., 1992), human diseases (Kazazian Jr, 1998), environmental adaptation (Casacuberta and González, 2013) and genome evolution (Kazazian, 2004). The evolutionary dynamics of TEs have been extensively studied, especially in the model organism *D. melanogaster* (Burt and Trivers, 2008). One particularly interesting feature that has emerged from these studies is that the vast majority of TE insertions in *D. melanogaster* tend to be at low population frequencies (Charlesworth et al., 1994, 1992; Sniegowski and Charlesworth, 1994; Biémont et al., 1994; Petrov et al., 2011; Montgomery and Langley, 1983; Kofler et al., 2012; Brookfield, 1986; Maumus et al., 2015). Recently, this pattern was also found in the closely related *D. simulans* (Kofler et al., 2014). Fixed TE insertions are largely restricted to low recombining regions (Bartolomé and Maside, 2004; Bartolomé et al., 2002; Kofler et al., 2012; Petrov et al., 2011) and to a few TE families (Kofler et al., 2012; Petrov et al., 2011, 2003;Hey, 1989). Two competing, but not mutually exclusive, models have been proposed to account for this predominance of low frequency insertions (Barrón et al., 2014). The transposition-selection balance model states that the abundance of most TE families is in an equilibrium, where the gain of novel insertions due to transposition equals the loss of copies by negative selection. (Charlesworth and Langley, 1989; Petrov et al., 2003, 2011; Nuzhdin, 1999; Blumenstiel et al., 2013). According to this model the low population frequency of TE insertions is mostly due to strong purifying selection acting against TE insertions. By contrast, the transposition burst model assumes that TE insertions with low frequencies are the consequences of recent bursts of TE activity (Bergman and Bensasson, 2007; Lerat et al., 2011; Koer et al., 2012).

A recent comparison of *D. simulans* and *D. melanogaster* identified substantial differences in TE abundance between these two species (Kofler et al., 2014). We proposed that these differences were probably due to an increased TE activity that could have been triggered by the recent habitat expansion of the two species (Kofler et al., 2014). If this hypothesis holds, endemic species are expected to have lower TE activities and thus fewer TE insertions with low frequencies.

We show that in contrast to *D. melanogaster* and *D. simulans* most most TE insertions in the island endemic *D. mauritiana* are fixed (66%). We propose that differences in the abundance of low frequency insertions between *D. mauritiana* (*f* ≤ 0.2, 18.0%) and *D. simulans* (64.3%), are likely due to different activities of up to 47 TE families. This suggests that activity of multiple TE families in *Drosophila* substantially changed over very short evolutionary time scales (<250,000 years), lending support to the transposition burst model of TE evolution. We propose that these differences in TE activity could be due to the different ecologies of the two species which may result in different opportunities for acquiring active TEs by horizontal transfer and different rates of loss of active TE families by vertical extinction.

## Results

Short read sequencing from pooled individuals [Pool-seq (Schlötterer et al., 2014)] has been shown to be an excellent approach to measure the population frequency of TE insertions on the genomic scale (Kofler et al., 2012, 2014; Kim et al., 2014). We compared TE abundance in a population of the island endemic *D. mauritiana* [data from Nolte et al. (2012)] to populations of the two cosmopolitan species *D. simulans* and *D. melanogaster* [data from Kofler et al. (2014); Nolte et al. (2012)]. The *D. simulans* and *D. melanogaster* populations were sampled in 2013 in Kanonkop (South Africa) (Kofler et al., 2014) and *D. mauritiana* was sampled between 2006 and 2009 from multiple locations in Mauritius (Nolte et al., 2012). We annotated TE insertions in all three reference genomes *de novo* (supplementary material and methods 4.2) as outlined in Kofler et al. (2014) and created a TE annotation for the *D. mauritiana* reference genome (Nolte et al., 2012). The TE abundance was estimated with PoPoolationTE (Kofler et al., 2012) after standardizing the physical coverage [numbers of paired-end reads spanning a TE insertion site (Meyerson et al., 2010)] to 60 in all populations. We only considered TE insertions in orthologous regions, i.e. regions present in the assemblies of all three species (see supplementary material and methods 4.2). This procedure permits a direct comparison of TE abundance between species. The impact of the various steps in our pipeline is detailed for every TE family in supplementary file 2.

### TE abundance in D. *mauritiana* and D. *simulans*

*D. mauritiana* contains significantly fewer TE insertions than *D. simulans (Dmau* = 2, 764, *Dsim* = 8, 056; Chi-square test, χ^2^ = 2, 588.3, *p < 2.2e –* 16) and this pattern is seen for all three TE orders (LTR *Dmau* = 532, *Dsim* = 1, 811; non-LTR *Dmau* = 404, *Dsim* = 1, 259; TIR *Dmau* = 1, 787, *Dsim* = 4, 737). Out of 7,097 *D. simulans* insertions for which population frequency estimates could be obtained (non-overlapping TE insertions) 1, 516 (21.4%) are fixed (f ≥ 0.9; allowing for some error) while in *D. mauritiana*, 1, 710 out of 2, 586 (66.1%) insertions are fixed (supplementary table 1). Despite the lower number of TE insertions *D. mauritiana* has more fixed insertions than *D. simulans (Dmau* = 1, 710, *Dsim* = 1,516, Chi-square test, χ^2^ = 11.7, *p* = 0.00063). This difference in fixed TE insertions is largely explained by a few TE families (*f* ≥ 0.9; top three in descending order *INE-1*: *Dmau* = 1,140, *Dsim* = 996; *roo*: *Dmau* = 88, *Dsim* = 67; *Crla*: *Dmau* = 43, *Dsim* = 32). The striking difference in overall copy numbers between the two species is mostly due to the about tenfold higher abundance of low frequency insertions in *D. simulans* (f ≤ 0.2; *Dmau* = 466 (18.0%), *Dsim* = 4, 562 (64.3%); χ^2^ = 3, 336.8, *p* < 2.2e – 16; supplementary fig. 1). This difference in low frequency insertions holds for all chromosome arms (fig. 1; supplementary table 1) and all TE orders (*f* ≤ 0.2; TIR *Dsim* = 2, 629, *Dmau* = 191, Chi-square test, χ^2^ = 2,107.7, *p* < 2.2e – 16; LTR *Dsim* = 1,147, *Dmau* = 153, Chi-square test, χ^2^ = 760.0, *p* < 2.2e – 16; non-LTR *Dsim* = 746, *Dmau* = 129, Chi-square test, χ^2^ = 435.1, *p* < 2.2e – 16). A more detailed analysis showed that 47 TE families had significantly fewer low frequency insertions in *D. mauritiana* (*f* ≤ 0.2; Chi-square test *p* ≤ 0.05; fig. 2), while only 3 families, including the intensely studied *Mariner* (Hartl et al., 1997; Lohe et al., 1995; Jacobson et al., 1986), had fewer low frequency insertions in *D. simulans* (fig. 2).

**Figure 1:**
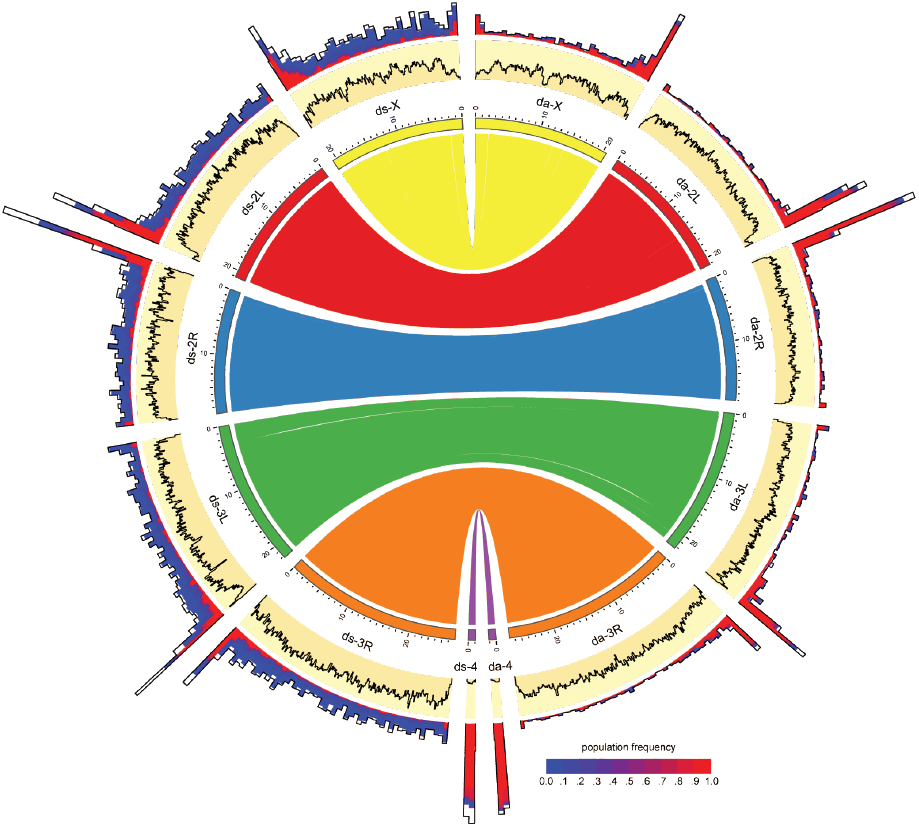
Distribution of TE insertions in a natural population of *D. simulans* (ds) and *D. mauritiana* (da). The TE distribution (outer graph) and the nucleotide polymorphism (Θ*π*, yellow inner graph) is shown. TE abundance is shown for 500kb windows, whereas the nucleotide diversity is shown for 100kb windows. For overlapping TE insertions (white) no estimates of population frequencies could be obtained. The relationship between the reference genomes is shown in the inside. The maximum nucleotide diversity of the plot is 0.0192 and the maximum number of TE insertions 275

**Figure 2:**
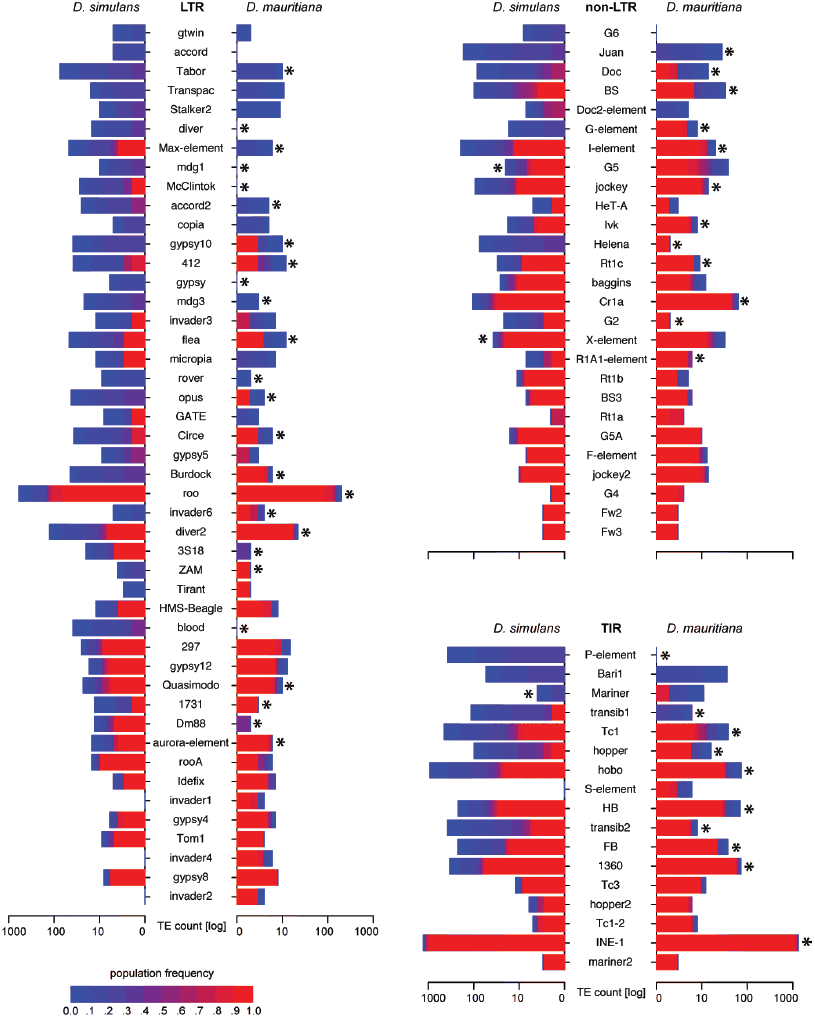
Abundance of different TE families in a natural *D. simulans* and *D. mauritiana* population. Only TE families with more than 10 insertions are shown. Significant differences in the abundance of low frequency insertions are indicated at the species having the lower counts (’*’; Chi-square test; *p* < 0.05). Foldback is grouped with TIR solely for graphic reasons.

### Robustness of the contrasting TE abundance pattern

Given the implicit challenges of cross-species comparisons, we carefully scrutinized our analysis to ensure that our result reflect a biological pattern rather than an artefact of the analysis: 1) Since the reported difference in low frequency insertions is consistently found in all steps of our pipeline (supplementary file 2) we can rule out that one or more filtering steps in the pipeline have caused this pattern. Even in the least processed data, 7.3% of the reads align to TE sequences in *D. simulans* while only 4.3% align to TEs in *D. mauritiana* (supplementary file 2). 2) Even without adjusting the physical coverage in both species, we find fewer TE insertions in *D. mauritiana*, despite this data set has a higher coverage (supplementary file 2; supplementary material and methods 4.1). 3) The *D. simulans* and *D. melanogaster* data were obtained from a single population but the *D. mauritiana* sample was composed of flies from multiple collections at different time points and locations (Nolte et al., 2012; Kofler et al., 2014). Although no population structure could be detected in *D. mauritiana* (Nunes et al., 2010), we tested if combining samples from different populations may cause a bias against low frequency TE insertions. We used an additional *D. simulans* population composed of flies sampled from multiple locations at different years (central Africa between 2001 and 2009; for an overview of all population see supplementary table 3). Although this *D. simulans* population has markedly fewer reads than the *D. mauritiana* population, which strongly favours identification of low frequency insertions in *D. mauritiana*, we still found a highly significant excess of low frequency insertions in *D. simulans* (*f* ≤ 0.2; *Dsim*_*ca*_ = 2, 911, *Dmau* = 466, χ^2^ = 1770.2 *p* < 2.2e – 16; supplementary file 2). 4.) It may be possible that a higher sequence divergence of TE insertions in *D. mauritiana* results in a smaller fraction of mapped TE reads and thus a lower abundance of TE insertions. Nevertheless, we consider this hypothesis unlikely since our pipeline takes sequence divergence into account by mapping reads to consensus TE sequences as well as to all diverged copies of a TE family found in a reference genome (using RepeatMasker and sensitive search settings). Furthermore, the same *de novo* annotation procedure, which relies on consensus TE sequences mostly derived from *D. melanogaster* (Quesneville et al., 2005), has been applied to both species. Since both species split after the divergence from *D. melanogaster* they are expected to have a similar divergence to *D. melanogaster* (Nolte et al., 2012). In agreement with this we detect no lineage specific TE families, i.e. we find more than 100 reads mapping to all TE families in both species [with two exceptions: the P-element is missing in *D. mauritiana* (Kofler et al., 2015) and *Stalker3* is absent in *D. simulans*; supplementary file 2]. Additionally, sequence divergence is expected to affect fixed TE insertions, which are enriched for old and inactive TE families (Kofler et al., 2014, 2012), more than low frequency insertions which are mostly young insertions derived from active copies that preserved functionality by escaping accumulation of mutations. In contrast to this expectation we found significantly more fixed TE insertions in *D. mauritiana* than in *D. simulans*, which suggests that higher sequence divergence of *D. mauritiana* TEs is not affecting our results. 5.) In contrast to the trend of a higher TE abundance in *D. simulans* in our data, Mariner insertions were previously shown to be more abundant in *D. mauritiana* (Jacobson et al., 1986; Hartl et al., 1997). Since our analyses confirm this pattern (*Dsim* = 4, *Dmau* =11; supplementary file 2), we conclude that our observation of a low TE abundance in *D. mauritiana* is not due to a general bias against identification of TE insertions in this species.

### Causes for the contrasting TE abundance pattern

Which evolutionary forces could be responsible for the parallel divergence of the number of low-frequency insertions across multiple TE families between two closely related species? In the following we discuss two, not mutually exclusive, hypotheses: 1) differential TE activity and 2) different selection efficacy. Since most TE insertions are deleterious (Burt and Trivers, 2008), differences in selection efficacy between species could cause the observed pattern. Both the population size (Charlesworth and Charlesworth, 1983; Kofler et al., 2014; González and Petrov, 2012) and the recombination rate (Dolgin and Charlesworth, 2008; Kofler et al., 2012) have frequently been shown to affect the efficacy of selection against TE insertions in natural populations. Since the efficacy of selection is higher in large populations (Hartl and Clark, 1997), the number of TE insertions, including low frequency insertions, should be lower in large populations (Kofler et al., 2014; González and Petrov, 2012). We used the nucleotide diversity *π* (Nei and Li, 1979) to compare the population size estimates in both species. The nucleotide diversity in *D. mauritiana* is lower than in *D. simulans* (average over all 100kb windows in orthologous regions; *π_Dmau_* = 0.0085, *π_Dsim_* = 0.0112; fig. 1) suggesting that *D. mauritiana* has a smaller population size than *D. simulans*, which is also consistent with the geographic distribution of the two species (Lachaise et al., 1988). However, a smaller population size as found in *D. mauritiana* could also lead to a loss of low frequency insertions by decreasing the efficacy of selection, thus allowing TEs to more rapidly fix (Lee and Langley, 2010). We consider it unlikely that this could be responsible for a reduced abundance of low frequency insertions in *D. mauritiana*: out of the 47 TE families with significantly fewer low frequency insertions in *D. mauritiana*, only 24 (51%) have more fixed insertions in *D. mauritiana* while the remaining 23 families (49%) either have equal or higher numbers of fixed insertions in *D. simulans*. Consequently it is unlikely that differences in the population size between the two species could account for the divergent abundance of low frequency insertions.

Alternatively a higher recombination rate could result in more ectopic recombination and less linkage between sites and therefore to an increased selection intensity against TE insertions (Dolgin and Charlesworth, 2008; Kofler et al., 2012). The genetic map of *D. mauritiana* is about 1.4 times (= 1.8/1.3) larger than the map of *D. simulans* (True et al., 1996). Assuming equal genome sizes for both species (Boulesteix et al., 2006), *D. mauritiana* should have an about 1.4 times higher recombination rate than *D. simulans*, suggesting that purifying selection against TE insertions is stronger in *D. mauritiana*. To test if recombination rate were influencing the abundance of low frequency insertions we mad use of recombination rate differences among chromosomes. True et al. (1996) reported that *D. mauritiana* chromosomes X, 2 and 3 have an about 1.8 (= 1.8/1.0), 1.23 (= 1.6/1.3) and 1.235 (= 2.1/1.7) fold larger genetic map than *D. simulans* chromosomes. Thus the X chromosome has the most pronounced differences in recombination rate between the two species and chromosome 2 the least. Despite these differences in recombination rates both chromosomes showed similar heterogeneity in the number of low frequency insertions between the two species (*f* ≤ 0.2, *Dmau*_*X*_ = 104, *Dsim*_*X*_ = 935, *Dmau*_*2*_ = 174, *Dsim*_*2*_ = 1,652, Fishers exact test, *p* = 0.6938). Since, X-chromosome and autosomes differ in many features other than recombination rates (Vicoso and Charlesworth, 2006), we further exploited local recombination rate heterogeneity on chromosome 3L (True et al., 1996). While the chromosome-wide recombination rate is higher in *D. mauritiana*, the local recombination rate between polytene bands 250 and 500 (measured from the centromere) on chromosome 3L is higher in *D. simulans* (True et al., 1996). Nevertheless, we still find fewer low frequency TE insertions in this genomic region in *D. mauritiana* (Chi-square test; *p* < 2.2e – 16; supplementary results 3.1). This suggests that recombination rate differences are not sufficient to explain the differences in TE composition between *D. simulans* and *D. mauritiana*.

Irrespective of whether selection efficacy is mediated by recombination rate or effective population size, differences will be reflected in the site frequency spectrum: stronger purifying selection results in a lower frequency of segregating TEs. To avoid misleading signals from ancestral insertions that occurred before the two species split, we only focussed on species specific TE insertions and compared the mean population frequencies. Interestingly, we found a higher mean population frequency of lineage specific TEs in *D. mauritiana* than in *D. simulans (Dmau* = 0.493, *Dsim* = 0.147, Wilcox rank sum test *W* = 3391131, *p* < 2.2e – 16). The higher population frequency of *D. mauritiana* insertions is consistent for all three TE orders (LTR *Dmau* = 0.419, *Dsim* = 0.153, Wilcox rank sum test *W* = 226682.5, *p* < 2.2e – 16; non-LTR *Dmau* = 0.341, *Dsim* = 0.156, Wilcox rank sum test *W* = 101537, *p* = 2.891e – 06; TIR *Dmau* = 0.606, *Dsim* = 0.144, Wilcox rank sum test *W* = 981319, *p* < 2.2e – 16; TIR without INE-1 *Dmau* = 0.264, *Dsim* = 0.106, Wilcox rank sum test *W* = 300776, *p* = 0.0042). Furthermore, 29 out of 39 TE families, with significantly different abundance of low frequency insertions between both species (fig. 1) and at least one lineage specific TE insertion in both species (39 out of 47), had on average a higher population frequency in *D. mauritiana* while 10 had a higher population frequency in *D. simulans* (supplementary file 3). The elevated population frequencies of *D. mauritiana* specific TE insertions persist when we exclude fixed insertions *f* ≥ 0.9; *Dmau* = 0.262, *Dsim* = 0.117, Wilcox rank sum test *W* = 1860001, *p* < 5.1e – 15).

Given that a range of different tests failed to provide convincing support for the hypothesis that the efficacy of selection against TE insertions explains the lower number of segregating TEs in *D. mauritiana* compared to *D. simulans*, an alternative explanation is required. We propose that the transposition activity differs between the two species, with the majority of families being more active in *D. simulans* (47 families). Nevertheless, 3 TE families, including the well-studied Mariner element (Hartl et al., 1997; Lohe et al., 1995; Jacobson et al., 1986), have more low frequency insertions in *D. mauritiana* and may be more active in this species (supplementary file 4).

### Comparison with *D. melanogaster*

It may be tempting to assume that low TE activity in *D. mauritiana* is a derived property, as *D. melanogaster* and *D. simulans* both have more low frequency insertions (f < 0.2; *Dmau* = 466 (18.0%), *Dmel* = 9,488 (81.7%), *Dsim* = 4,562 (64.3%), supplementary file 4) and consequently may have more active TEs. We caution, however, that this interpretation is too simplistic, since the rapid activity change between *D. mauritiana* and *D. simulans* suggests that several changes in TE activity may have occurred since the split of *D. melanogaster* and the *D. simulans* group. Despite *D. simulans* and *D. melanogaster* sharing a high TE activity, the profile of active TE families in *D. simulans* is more similar to *D. mauritiana* than to *D. melanogaster* (Spearman correlation of the abundance of low frequency insertions, *f* < 0.2, for all TE families; Dmau-Dsim *ρ* = 0.58, *ρ* = 8.3e – 11; Dsim-Dmel *ρ* = 0.43, *ρ* = 4.9e – 06; supplementary file 4).

## Discussion

In this report we compare for the first time the genomic distribution of TE insertions in three closely related *Drosophila* species on a population scale. By standardizing the Pool-Seq data to the same physical coverage and using an identical pipeline for TE identification in all three species we minimize potential biases of interspecific comparisons. While the TE landscape of *D. simulans* and *D. melanogaster* populations fit the previously described predominance of low frequency TE insertions (Charlesworth et al., 1994, 1992; Sniegowski and Charlesworth, 1994; Biémont et al., 1994; Petrov et al., 2011; Montgomery and Langley, 1983; Kofler et al., 2012, 2014), in *D. mauritiana*, the pattern is fundamentally different. We show that the island endemic *D. mauritiana* not only has fewer TE insertions than the other two species, but the insertions have a significantly higher population frequency, with the majority of them being fixed. This unexpected TE distribution could be explained either by stronger purifying selection in *D. mauritiana*, removing novel insertions, or a higher transposition rate in *D. simulans* and *D. melanogaster*. We carefully scrutinized the *D. mauritiana* data for any signals of higher selection efficacy, but did not detect support for this hypothesis. Therefore, we concluded that transposition rate heterogeneity is the most likely explanation for the contrasting TE distribution between the species. Such rapid changes in TE activity affecting a broad range of TE families has only previously been reported in plants. One particular impressive example is the explosive activity of 11 LTR families in maize which led to a doubling of the genome size within 3 million years (SanMiguel et al., 1998). No marked differences in genome size were, however, reported for *D. simulans* and *D. mauritiana* (Boulesteix et al., 2006), probably because the higher TE activity in *D. simulans* is mostly reflected in a high abundance of low frequency insertions, which have on the average a small impact on genome size. We do not consider it very likely that a similar genome expansion will be seen in *D. simulans* since the large effective population size allows for a very efficient selection against deleterious TE insertions.

With the three *Drosophila* species being closely related and sharing almost all TE families, it is very unlikely that simple structural differences could be responsible for the divergence in transposition rates. Rather, we propose that two different, but not mutually exclusive, processes related to the contrasting ecology of the species may be responsible for the heterogeneity in transposition rates. First, environmental stress may activate TEs (Capy et al., 2000) and colonizing species, like *D. simulans* and *D. melanogaster* (Lachaise et al., 1988), may be exposed to more environmental stress than species that remained in the ancestral habitat such as *D. mauritiana* (Lachaise et al., 1988). So far, stress was only shown to activate a few TE families, like 412 and hobo (Capy et al., 2000), and therefore it remains unclear if stress can account for the observed activation of 47 TE families. Second, the balance between the two opposing forces of vertical extinction and horizontal transmission may be shifted between *D. simulans* and *D. mauritiana*. Vertical extinction, i.e. the loss of active TE copies, may result from competition between TE families, the accumulation of deleterious mutations and the evolution of host repression of TEs (Burt and Trivers, 2008). On the other hand, active TE copies may be gained by horizontal transmission from different species (Bartolomé et al., 2009; Sánchez-Gracia et al., 2005; Silva et al., 2004). The number of active TE copies segregating in a population may thus be the outcome of these two opposing forces. Using a simple model Kaplan et al. (1985) showed that vertical extinction may be rapid in small populations, and we found that *D. mauritiana* likely has a smaller population size than *D. simulans*. Furthermore, the colonizing *D. simulans* may have had more opportunities for acquiring active TEs by HT than the island endemic *D. mauritiana*, especially given that opportunities for HT increase with population size and species diversity in the habitat, both of which may be low for island endemic species (MacArthur and Wilson, 1967). Reduced opportunity for HT has also been suggested as explanation for the low TE content of the island endemic *D. grimshawi* (Drosophila 12 Genomes Consortium, 2007) and could also be responsible for the high fraction of fixed insertions in the endemic D. algonquin (Hey, 1989). It is therefore possible that vertical extinction of TE families predominates in *D. mauritiana* while horizontal acquisition of active TE copies is more frequent in *D. simulans*. The presence of all TE families (except P-element and Stalker3) in the three species may be interpreted to counter this hypothesis, which requires some horizontal transfer of active TEs, as HT is expected to cause a patchy distribution of TEs in the phylogeny of species (Schaack et al., 2010; Loreto et al., 2008; Silva et al., 2004). However, abundant HT, as for example found for the P-element which invaded two *Drosophila* species within one century (Kofler et al., 2015), could also lead to the presence of all TE families in the three species.

Our results also have bearing on a long-standing debate about the evolutionary dynamics of TEs (Barrón et al., 2014). The transposition-selection balance model assumes that the abundance of most TE families reflects the equilibrium of gains of novel insertions by transpositions and the loss of copies by negative selection. (Charlesworth and Langley, 1989; Petrov et al., 2003, 2011; Nuzhdin, 1999). Thus low population frequencies of TE insertions are the outcome of strong purifying selection acting against TE insertions. By contrast, according to the transposition burst model, TE insertions with low frequencies are the consequence of recent increase in TE activity (Kofler et al., 2012; Bergman and Bensasson, 2007; Lerat et al., 2011; Blumenstiel et al., 2013). Our finding of substantial differences in activity of multiple TE families between two closely related species at short time scales raises the important questions of whether the TE distribution in *D. simulans* and *D. melanogaster* has already reached an equilibrium state. Assuming that habitat expansions and stressful environments modulate TE activity, it appears possible that the distribution of TEs rarely reaches an equilibrium state. We anticipate that future work analyzing multiple populations of related species with different ecologies may shed further light on the forces shaping the evolutionary dynamics of TEs.

## Material and Methods

We measured TE abundance in one population of *D. mauritiana*, two populations of *D. simulans* and one population of *D. melanogaster* using previously published Pool-seq data (Nolte et al., 2012; Kofler et al., 2014). For details about the samples used in this study see supplementary table 4.1. We *de novo* annotated TE insertions in the reference genomes of *D. simulans* (r1.0 Palmieri et al., 2014), *D. mauritiana* (r1.0 Nolte et al., 2012) and *D. melanogaster* (v6.03; dos Santos et al., 2015) and identified TE insertions with PoPoolationTE (Kofler et al., 2012). In contrast to our previous work (Kofler et al., 2014) we included the canonical sequence of *Mariner,* which was discovered in *D. mauritiana* (Hartl et al., 1997), into our pipeline for estimating TE abundance. Pairwise nucleotide diversity was estimated for a natural population of *D. mauritiana* (Nolte et al., 2012) and a natural population of *D. simulans* from South Africa (Kofler et al., 2014) using PoPoolation (Kofler et al., 2011). Orthologous regions between *D. simulans*, *D. melanogaster* and *D. mauritiana*, i.e. regions occurring in the assemblies of all three species, were identified with MUMmer (v3.23; nucmer) (Kurtz et al., 2004). TE insertions at similar genomic positions in *D. mauritiana* and *D. simulans* were identified by reciprocally aligning 1000 bp regions flanking each TE insertion to the respective reference genomes with bwa (v0.7.5a) (Li and Durbin, 2010) and scanning for insertions of the same family within these boundaries. All statistical analysis was done with R (R Core Team, 2012). For details see supplementary material and methods. The TE annotation of the *D. mauritiana* genome and the TE abundance in the *D. mauritiana* population have been made publicly available (https://sourceforge.net/p/popoolationte/wiki/pdmau/).

## Acknowledgments

We thank all members of the Institute of Population Genetics for feedback and support. This work was supported by the ERC grant Archadapt.

## Supplementary files

- **Supplementary file 1** Supplementary figures, tables and material and methods (pdf)
- **Supplementary file 2** A table containing for every TE family detailed statistics about the number of mapped reads, paired end fragments supporting a TE insertions and TE insertions identified with PoPoolationTE (xlsx)
- **Supplementary file 3** A table containing for every TE family the number of lineage specific TE insertions (xlsx)
- **Supplementary file 4** A table containing for every TE family the number of low frequency insertions (*f* ≤ 0.2) (xlsx)

